# CACNA1C TS-II variants alter single-cell dynamics in computational models of cortical pyramidal cells

**DOI:** 10.1101/2025.07.02.662758

**Authors:** M. Moore, M-L. Linne, T. Mäki-Marttunen

## Abstract

Timothy Syndrome (TS) is a rare multi-system disorder and a monogenic calcium channelopathy. Previous computational work on this disorder has focused on the myocardium, ignoring the effects of TS on neural development and its strong association with autism spectrum disorder. Variation in the TS-causative gene, *CACNA1C*, is indeed also associated with a variety of complex neurodevelopmental and neuropsychiatric disorders. We apply computational methods, drawing on experimental data, to understand the mechanisms of calcium dysregulation in TS, and validate our findings in four well-established multi-compartmental neuronal models. *CACNA1C* encodes the L-type voltage-gated calcium channel, Ca_v_1.2, which modulates neuronal excitability and several activity-dependent pathways. We investigate two mutations in *CACNA1C* causative of TS type II, Ca_v_1.2^G406R^ and Ca_v_1.2^G402S^. Both variants show a loss of voltage-dependent inactivation and changes in their voltage dependence of activation. We incorporate the altered steady-state activity of these variants with additional morphological data indicating a significant increase in activity-dependent dendritic retraction of layer II/III pyramidal cells in TS mutant neurones. Our findings replicate experimental work suggesting that increased Ca^2+^ flux reduces firing frequency but does not affect the rheobase current for action potential initiation. Furthermore, models expressing dendritic Ca_v_1.2 current show altered apical-somatic signal integration in mutant neurones. All models with shortened dendrite morphology show hyperexcitability, denoted by reduced rheobase current and increased responsiveness to current injection compared to their full-length counterparts. Importantly, our approach identifies robust and testable predictions on the impacts of TS on single-cell dynamics. We also discuss the broader implications of our findings for other Ca^2+^-related neurodevelopmental and neuropsychiatric disorders.

## 1 Introduction

The disruption of calcium ion (Ca^2+^) influx and the consequent changes in neuronal excitability are a common factor in many neuropsychiatric and neurodevelopmental disorders [1–7]. In particular, *CACNA1C*, the gene that encodes the *α* subunit of the L-type Ca^2+^ channel, Ca_v_1.2, is among the most robustly linked psychiatric risk genes from multiple genome-wide association meta-analyses [2, 8–12]. However, numerous other genes associated with neuronal function are implicated in the pathophysiology of neuropsychiatric disorders, making it difficult to discern the unique consequences of *CACNA1C* alterations on neuronal activity [13]. A dominant mutation in *CACNA1C* is also the cause of the rare multi-system disorder, Timothy Syndrome (TS) [12, 14]. *CACNA1C* is highly expressed in brain and cardiac tissue [15]. TS patients universally present with cardiac arrhythmias, and many also exhibit neurodevelopmental delay and autism spectrum disorder symptomatology [16–18]. Although the channelopathy associated with TS is almost exclusively investigated in terms of heart failure, its monogenic cause removes much of the uncontrolled genetic variation that impedes neuropsychiatric research. Therefore, we propose it as a promising model to investigate the pathophysiology of Ca_v_1.2-related neuropsychiatric and neurodevelopmental disorders.

TS is the most penetrant monogenic form of autism spectrum disorder, with 60–80% of typical TS patients meeting the criteria [14]. In TS, a missense mutation induces prolonged Ca_v_1.2 channel activation and open probability, resulting in massive increases in Ca^2+^ influx [14, 16]. Although the electrophysiological effects and channel dynamics have been widely studied in TS (for a review of current knowledge, see [19]), the impact of TS mutations on single-neurone excitability remains largely unknown [20]. This limitation hampers our understanding of the altered cellular and circuit functions that underlie the pathology of TS and other types of autism spectrum disorder.

The current study aims to assess the effect of TS-associated Ca^2+^ channel alterations on cellular electrophysiology and network activity using computational modelling. We develop and compare four well-established biophysically detailed multi-compartmental neurone models and include parameters derived from *in vitro* and *in vivo* TS experiments. In particular, we use electrophysiological data associated with a variant of TS, TS-II, which manifests a more severe phenotype than TS-I [16]. We also employ morphological data indicating a significant activity-dependent dendritic retraction of layer II/III pyramidal cells (L23PCs) in *in vitro* human and *in vivo* murine models of TS [21, 22]. We integrate these data to investigate changes in single-cell electrophysiology caused by TS-II Ca_v_1.2 *α*-subunit variants. We find that 1) TS-II variants reduce firing frequency due to heightened Ca^2+^ influx, 2) shortened cells show increased excitability, and 3) these effects independently, and in combination, alter signal processing in the single cell. Our results provide a base computational unit from which to investigate network effects in TS and bridge the gap between genetic aetiology and behavioural characteristics. Furthermore, understanding the effects of dysregulated calcium dynamics in TS will help to elucidate the pathophysiology of more complex calcium-dependent neuropsychiatric disorders, particularly the highly comorbid autism spectrum disorder.

## 2 Methods

### Pyramidal Cell Models

We interrogated four well-established multi-compartmental models of cortical pyramidal cells (PCs) to determine the effects of TS variants on single-cell dynamics. TS-induced dendritic shortening was observed in L23PCs [21]. Hence, we compared a biophysically detailed L23PC model, obtained from the Blue Brain Project (BBP), to three alternative models. Our alternative models comprise two additional L23PC models [23, 24] with reduced current sets and one layer V (L5PC) model [25]. The final alternative model was selected as one of the best-validated multi-compartmental models in the field and simulates a full set of comparable ionic currents found in the BBP L23PC model (Table 1). Despite known interlayer morphological and biophysical differences between PC subtypes [26], this model provides a useful point of comparison.

**Table 1:** Origin of current equations in each model.

| Current | BBP | Branco | Mainen | Hay |
| --- | --- | --- | --- | --- |
| $I_{CaHVA}$ | Reuveni et al. [32] | Reuveni et al. [32] | Reuveni et al. [32] | Reuveni et al. [32] |
| $I_{CaLVA}$ | Avery and Johnston [33] | Destexhe and Huguenard [34] | | Avery and Johnston [33] |
| $I_{SK}$ | Köhler et al. [35] | Reuveni et al. [32] | Reuveni et al. [32] | Köhler et al. [35] |
| $I_{Kp}$ | Korngreen and Sakmann [36] | Hamill et al. [37] | Hamill et al. [37] | Korngreen and Sakmann [36] |
| $I_{Kt}$ | Korngreen and Sakmann [36] | | | Korngreen and Sakmann [36] |
| $I_{Kv3.1}$ | Rettig et al. [38] | | | Rettig et al. [38] |
| $I_m$ | Adams et al. [39] | Mainen and Sejnowski [24] | Mainen and Sejnowski [24] | Adams et al. [39] |
| $I_{Nap}$ | Magistretti and Alonso [40] | Huguenard et al. [41] | Huguenard et al. [41] | Magistretti and Alonso [40] |
| $I_{Nat}$ | Colbert and Pan [42] | | | Colbert and Pan [42] |

The BBP model, a reconstructed rat somatosensory L23PC (Figure 1), contains twelve ionic currents simulating passive leak currents (I*_leak_*); high and low voltage-activated Ca^2+^ currents (I*_CaHV A_* and I*_CaLV A_*); fast inactivating and persistent sodium (Na^+^) currents (I*_Nat_* and I*_Nap_*); and five potassium (K^+^) currents, including slow inactivating (I*_Kp_*), fast inactivating (I*_Kt_*), fast non-inactivating (I*_Kv_* _3.1_), muscarinic (I*_m_*), and small-conductance (I*_SK_*), for descriptions see Hay et al. [25]. We used the default version of the model, where the distribution of current densities were calculated to best reproduce the electrophysiological characteristics of the cell type [27, 28], and the reconstructed axon is replaced by a short artificial axon [27]. The model was obtained from the *BBP Neocortical Microcircuit Collaboration Portal*, for model files see: https://bbp.epfl.ch/nmc-portal/microcircuit.html#/metype/L23_PC_cADpyr/details.

**Figure 1:**
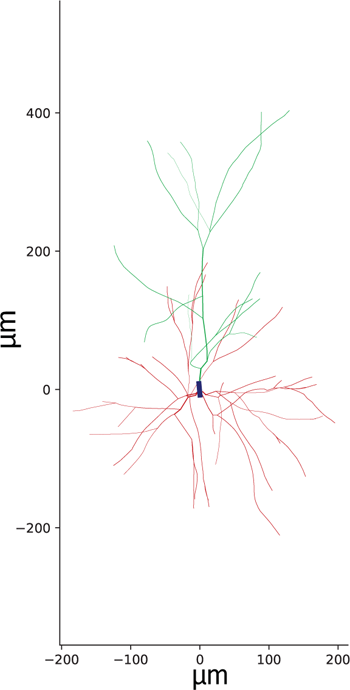
Reconstructed 2D morphology of a BBP L23PC model depicting the soma (dark blue), apical dendrites (green), and basal dendrites (red). Axon not shown.

Alternative L23PC models from Mainen and Sejnowski [24] (later referred to as the “Mainen model”) and Branco et al. [23] (henceforth the “Branco model”) have been used as a comparison. These models contain a reduced set of ion currents, comprising I*_CaHV A_*, I*_m_*, a non-specific Ca^2+^-dependent K^+^ current (I*_nsKCa_*), one voltage-dependent K^+^ current (I*_Kv_*), one voltage-dependent Na^+^ current and I*_CaLV A_*). The Branco model also contains I*_CaLV A_* (Table 1). A fourth model, taken from Hay et al. [25] (“Hay”), is a L5PC model containing 12 currents of the same origin as the BBP model (Table 1).

Crucially, unlike the BBP model, all three alternative models express I_CaHVA_-densities in dendritic compartments. This is in agreement with experimental data suggesting Ca_v_1.2 channels are present in the proximal and distal dendrites of L23 and L5 PCs [29–31].

### L-type Ca**^2+^** Channel Dynamics

High voltage activated (HVA) Ca^2+^ channels in the brain comprise all Ca*_v_*1 and Ca*_v_*2 subtypes [43], with Ca*_v_*1.2 being particularly highly expressed [44]. There is high similarity in the structure and binding sites of L-type Ca^2+^ channels, Ca_v_1.2 and Ca*_v_*1.3, that preclude *in vitro* investigations of single subtype conductance in neurones, therefore, there is little data available to estimate isolated Ca_v_1.2 current for *in silico* studies [15]. However, the available evidence suggests that Ca*_v_*1.3 accounts for just 10% of L-type Ca^2+^ channel expression in the brain and 15% of L-type Ca^2+^ channel current in the somatosensory cortex, with other L-type Ca^2+^ channels contributing minimally [44, 45]. Therefore, in this work we consider the I_CaHVA_ current to be Ca_v_1.2 current.

### TS-II L-type Ca**^2+^** Channel Dynamics

Among voltage-gated Ca^2+^ channels, L-type channels are characterised by large single-channel conductance, a high half-activation voltage of voltage-dependent activation (VDA) and slow voltage-dependent inactivation (VDI) [43]. TS mutations impact both the VDA and VDI of Ca_v_1.2. The two typical forms of TS, TS-I and TS-II, are caused by substitution mutations in the mutually exclusive splice variants exon 8a (E8a) or exon 8 (E8), respectively [14, 16]. The two splice variants are highly homologous but exhibit differences in temporal expression profiles and tissue-specific expression during adulthood [46]. As a result of the higher relative expression of E8 in brain and cardiac tissue, TS-II manifests as a more severe phenotype [16]. TS-II can be caused by a Glycine to Arginine substitution, G406R, or a Glycine to Serine substitution, G402S, in transmembrane segment 6 of domain 1 (S6-D1) of *CACNA1C* [16] (Figure 2). Notably, Ca_v_1.2^G402S^ patients typically show normal neurological development, while Ca_v_1.2^G406R^ patients present with the full neurological symptoms of TS, including autism spectrum disorder [16, 17, 47].

**Figure 2:**
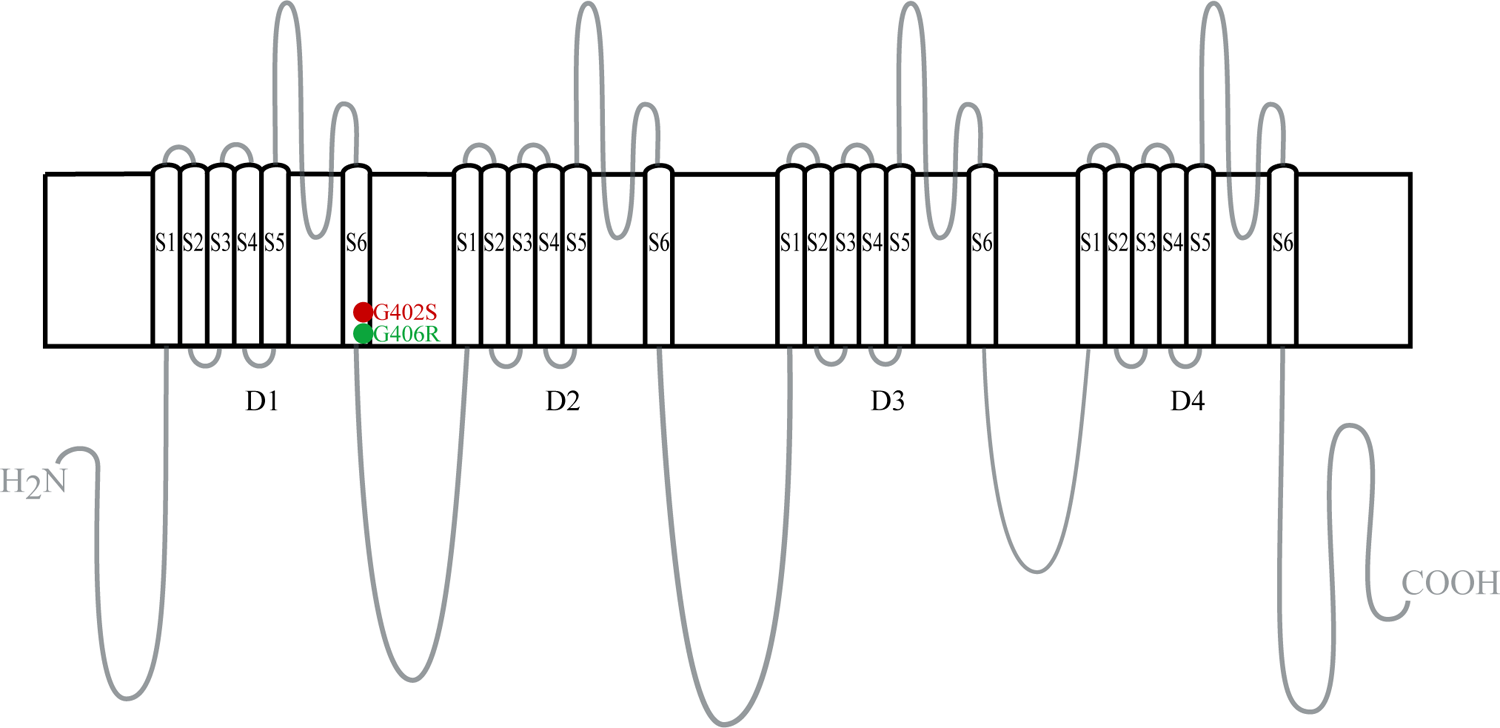
Ca_v_1.2 *α*-subunit in the membrane. The channel consists of four domains (D1-D4), each comprising six transmembrane segments (S1-S6). Red and green circles indicate the approximate locations of the TS-II substitution mutations G402S and G406R, respectively ([16]. These mutations occur in both Exon 8 and Exon 8a. G, Glycine; S, Serine; R, Arginine

TS-II mutations, Ca_v_1.2^G402S^ and Ca_v_1.2^G406R^, induce near complete loss of VDI [14, 16]. To incorporate this, an additional parameter, *h-baseline* (*h_bl_*), was added to the *h_∞_* inactivation curve equation as follows:

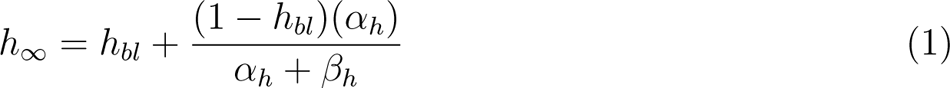

This creates a baseline value to which channel inactivation is limited. Ca_v_1.2^G402S^ and Ca_v_1.2^G406R^ channels have *h_bl_* values of 0.91 and 0.88, respectively, while wild-type (Ca_v_1.2^WT^) channels have a *h_bl_* of 0.0 (Table 2). Additionally, the Ca_v_1.2^G406R^ mutation causes a left-ward shift in the VDA curve by *−*9mV, whereas the Ca_v_1.2^G402S^ mutation causes an opposite shift in VDA by 2mV [16]. We model this by altering the half-activation voltage parameters of the models using the associated voltage-shift (*V*_shift_) value (Table 2, Figure 3).

**Figure 3:**
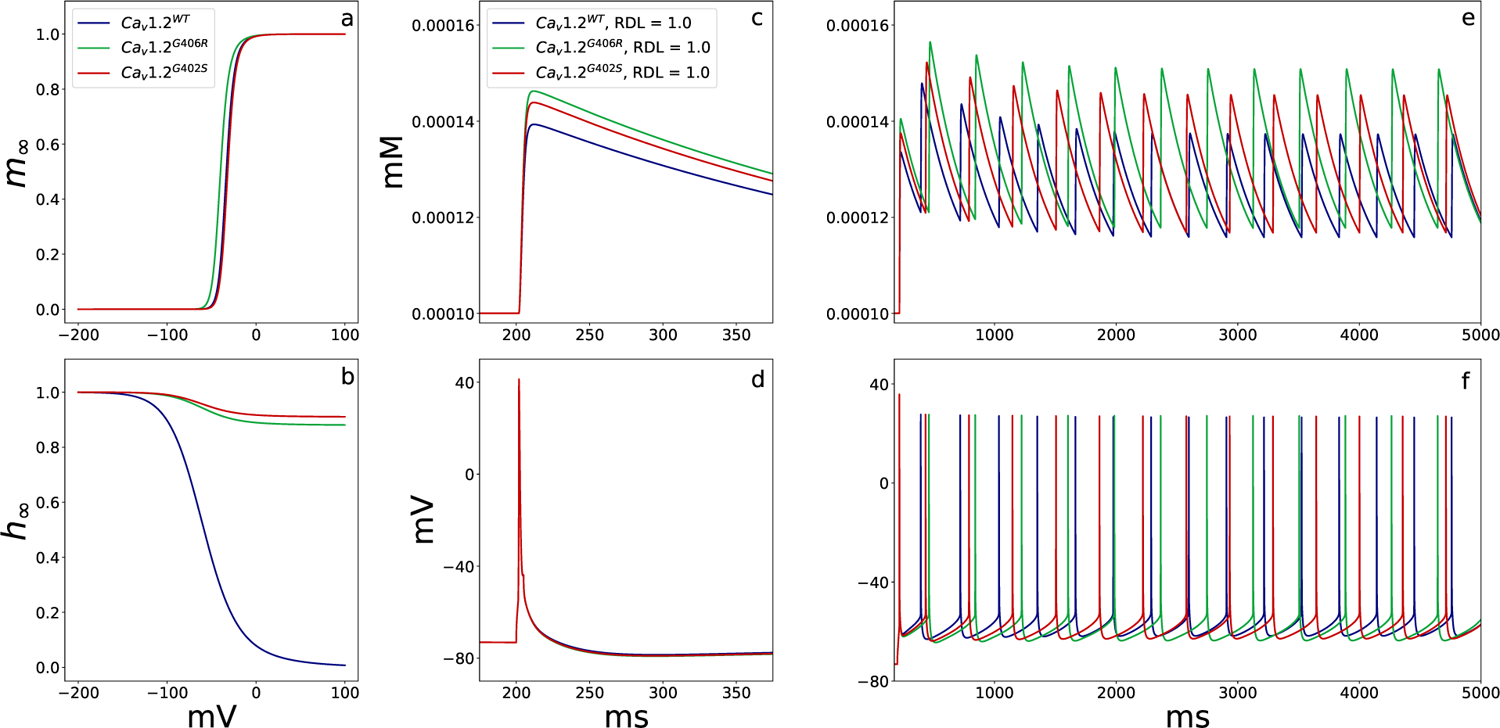
Altered gating kinetics of TS-II Ca_v_1.2 variants of the BBP model. Steady-state activation (a) and inactivation (b) curves for all channel variants. The Ca^2+^ response (c,e) and spiking behaviour (d,f) of WT and TS-II variant cells during short somatic square pulse stimulation (c-d) and long somatic square pulse stimulation (e-f). For the short stimulus, a 1.0nA short square pulse was applied for 5ms. For the long stimulus, a 0.2nA long square pulse was applied for 7800ms. Both stimuli were applied after a 200ms delay to allow the cell to reach equilibrium. Somal *V_m_* or Ca^2+^ concentration was then plotted against time. Here, the effects of the mutation-associated alterations of SSA and SSI without the effects on relative dendritic length (RDL) were considered.

**Table 2:**
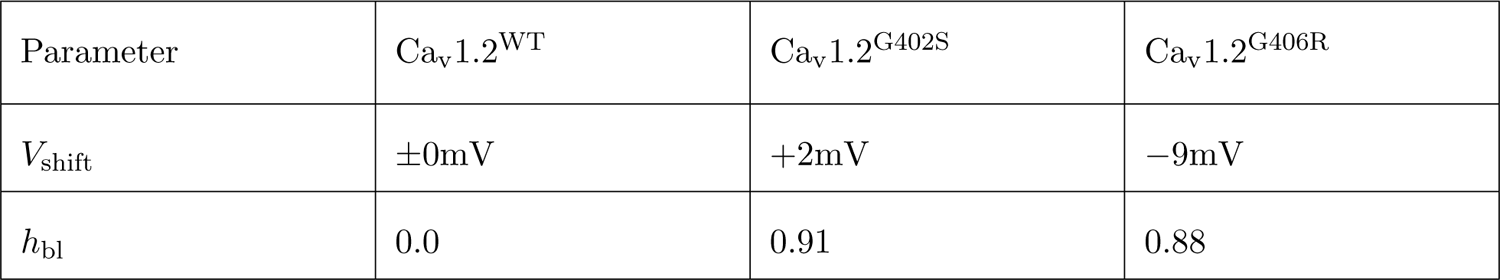
Voltage dependence of inactivation (VDI) and voltage dependence of activation (VDA) parameters altered in modelled TS-II variants.

TS mutations are autosomal dominant and homozygosity is non-viable [48, 49]. To model the heterozygosity, we created a second I_CaHVA_ current, I_CaHVA_TS, containing the single channel current data from [16] and replaced 50% of the total I_CaHVA_ density with this current. I_CaHVA_TS density was further reduced to account for the splicing pattern of CACNA1C E8 and E8a. The unaffected E8a splice variant is expressed in 23% of neuronal Ca_v_1.2 channels [14]. Therefore, in TS-II, 38.5% of neuronal channels contain the substitution mutation.

### Dendritic Length

TS mutations also disrupt activity-dependent regulation of basal dendrite morphology, resulting in a reported 30% average reduction in length [21]. Although differences in the regulation of apical and basal dendrites exist, similarities in their development indicate similar molecular processes are involved in extension and retraction [50–52]. As such, we hypothesised that a similar truncation would occur in both both basal and apical dendrites. This was modelled by reducing the relative length of each apical and basal dendritic compartment, after model initiation.

### Simulation of Electrophysiological Responses

All models were adapted for use as TS-II model neurones by adjusting HVA Ca^2+^ current parameters and dendrite length, as described above. Simulation protocols were created to assess spiking and signal integration behaviours. All protocols were initiated after a 200ms delay to allow the cells to reach equilibrium. Stimuli were applied to either the somatic compartment or an apical location 140–200*µ*m from the soma. When stimulated at the apical dendrite, the stimulus location was always relative to the length of the dendrite.

A short square pulse of 1.0nA for 5ms and a long square pulse of 0.2nA for 7800ms were used to assess action potential and Ca^2+^ dynamics in control and mutant BBP model neurones. Current step protocols for analysing the frequency-current relationship used the long square pulse with amplitudes ranging from 0–0.6nA in 0.01nA steps. Rheobase values, defined as the minimum current required to elicit an action potential, were determined using a binary search algorithm over 20 iterations with a 3800ms square pulse, with current values ranging from 0.0nA to 2.0nA. To investigate altered signal integration in TS neurones, an interstimulus interval protocol was adapted from Hay et al. [25] and Mäki-Marttunen et al. [53]. Two stimuli were applied to the neurone; a short square pulse (5ms) at the soma, and an excitatory postsynaptic potential pulse applied at a point 50–65% along the longest apical branch. The somatic stimulus was applied at 1000ms, and the order and interval between that and the apical stimulus were dictated by the inter-stimulus interval, which ranged from *−*20–40ms. For each model, the somatic and apical input amplitude was determined using aca binary search to find the minimum amplitude between 0–2nA that elicited at least one spike across a range of intervals, then increased by 5%. When assessing firing frequency and rheobase values, a spike recognition filter was applied with a repolarisation and depolarisation threshold of *−*10mv.

All simulations were run with NEURON 8.2 and Python 3.9.11. Simulation scripts are available at https://modeldb.science/2016667.

## 3 Results

### TS Variants Increase Intracellular Calcium Response to Stimulus

We implemented the steady-state changes found in Ca_v_1.2^TSII^ channels in Splawski et al. [16]. The adjusted half-activation voltage values – depending on the current origins (Table 2) – implemented the appropriate shifts in the steady-state activation (SSA) curves of Ca_v_1.2^G402S^ and Ca_v_1.2^G406R^ channels (Figure 3a). Similarly, introducing the VDI parameter, *h_bl_*, to adjust the baseline plateau of I_CaHVA_TS to 0.88 and 0.91 for Ca_v_1.2 and Ca_v_1.2, respectively (Table 2), achieved the drastic loss of steady-state inactivation (SSI) characterised in Splawski et al. [16] (Figure 3b). Due to adjustment for heterozygosity and the ratio of E8:E8a expression in the brain, only 38.5% of the total Ca*^HV^ ^A^*current was modelled with these kinetics, the remaining was modelled as WT current.

These altered kinetics cause an increase in the Ca^2+^ response to short and long stimulus durations (Figure 3c,e). This has no effect on the timing of the initial spike (Figure 3d) but during sustained depolarisation causes an elongation of the inter-spike interval (Figure 3f). This elongation happens because increased Ca^2+^ response induces larger SK channel activation, and increased SK channel activation results in a greater efflux of K^+^ and a prolonged afterhyperpolarisation phase (AHP) [54]. The effect of the TS variants are greater in Ca_v_1.2^G406R^, which is consistent with the reduced VDA increasing the probability of channels opening at lower *V* and resulting in greater Ca^2+^ influx.

### TS Variants Alter the Frequency-Current Relationship

We then assessed whether the increased Ca^2+^ current through TS variants would alter the frequency-current relationship (f-I) of the cellular response. In all models, TS variant SSA/SSI alterations affects firing frequency under long-duration stimulation (Figure 4) with-out impacting the rheobase–the minimum current required to initiate an action potential–for short-duration stimuli (Figure 4 inset axes). In the BBP model, and Mainen and Hay alternative models, the frequency-current (f-I) curve is flattened and firing frequency reduced. f-I curve flattening in response to increased Ca_v_1.2 activity and Ca^2+^ concentration is expected, and results from Ca^2+^-dependent activation of I*_SK_*currents [53, 55]. The Branco model shows an opposite trend, with the Ca_v_1.2^G406R^ variant causing increased cellular excitability, as denoted by the leftward shift of the curve (Figure 4b,f). This could be due to a reduced role for the I*_KCa_* in Branco, and as such the increased Ca^2+^ current is not counteracted by membrane hyperpolarisation.

**Figure 4:**
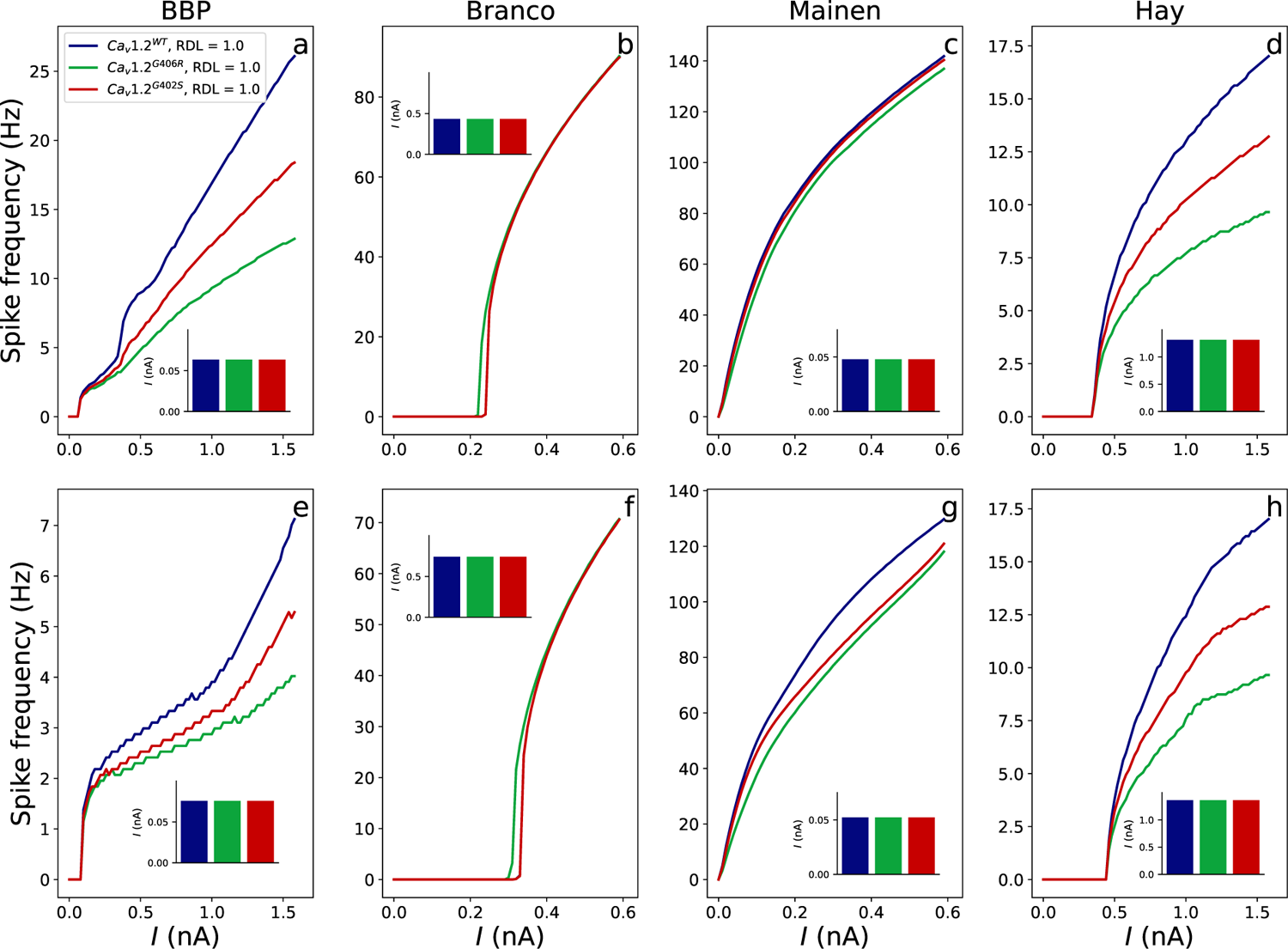
Impacts of the SSA/SSI alterations of TS variants on f-I curves and initial rheobase (inset axes) of multicompartmental L3PC models BBP (a,e), Branco (b,f), Mainen (c,g), and an L5PC Hay model (d,h). The f-I curves were created using a total simulation time of 10000ms. Current was applied for 7800ms, 200ms after initiation of the simulation, amplitude ranged from 0–0.6nA in 0.01nA steps. Spike frequency was calculated over the interval of 500–10000ms. A spike recognition filter was applied such that spikes were recognised only if the previous repolarisation valley dropped below *−*10mV and depolarisation peaked passed *−*10mV. Rheobase values were calculated based on 20 iterations of the bisection method. The stimulus was applied at somatic or apical (200*µ*m from the soma) locations. Upper panels (a-d) show the somatic stimulation paradigm and lower panels (e-h) show apical stimulation. The BBP and Hay models have a lower baseline firing frequency, as such a wider range of current amplitudes was used to ensure comparable firing rate scales with the other models. RDL, relative dendritic length.

For all models, the more extreme Ca_v_1.2^G406R^ variant produces a larger effect. These findings also concur with the known consequences to Ca^2+^ current of each variant, with the Ca_v_1.2^G406R^ variant causing greater Ca^2+^ influx and showing a more extreme effect on the f-I curve. These effects are seen for both somatic (Figure 4a-d) and apical stimuli (Figure 4e-h). The Mainen model, and to a lesser extent the Branco and Hay models, show more extreme effects when stimulated in the apical dendrite. This may be due to the presence of I_CaHVA_ current in the dendrites of these cells, heightening the effect of Ca_v_1.2 SSA/SSI alterations on cell activity.

The rheobase of all models are unaffected by SSA and SSI changes alone (Figure 4 inset axes). Physiologically, rheobase is unaffected by I*_SK_* currents due to the delayed activation of SK channels. Taken together, our data suggests that the altered gating kinetics of TS-II variants reduce firing frequency, and may reduce the threshold for repetitive firing, but have no effect on short-pulse rheobase in PCs.

### TS-Related Cellular Truncation Affect Spiking Behaviour

Dendritic shortening increases cellular excitation by reducing leak to the dendrites, causing a reduction of firing threshold and an increase in the firing frequency. The BBP model with TS-associated dendrite trunction shows a slight leftward f-I shift and drastically increased frequencies at high input currents (Figure 5a). The BBP and Branco models are particularly affected when stimulated at the apical dendrite location, at a distance relative to dendritic shortening: 200*µ*m from soma in WT and 140*µ*m from soma in TS-II variant cells. All three alternative models show either a leftward shift of the curve (Figure 5b,d) or an increased f-I gradient in response to both stimulation conditions (Figure 5c).

**Figure 5:**
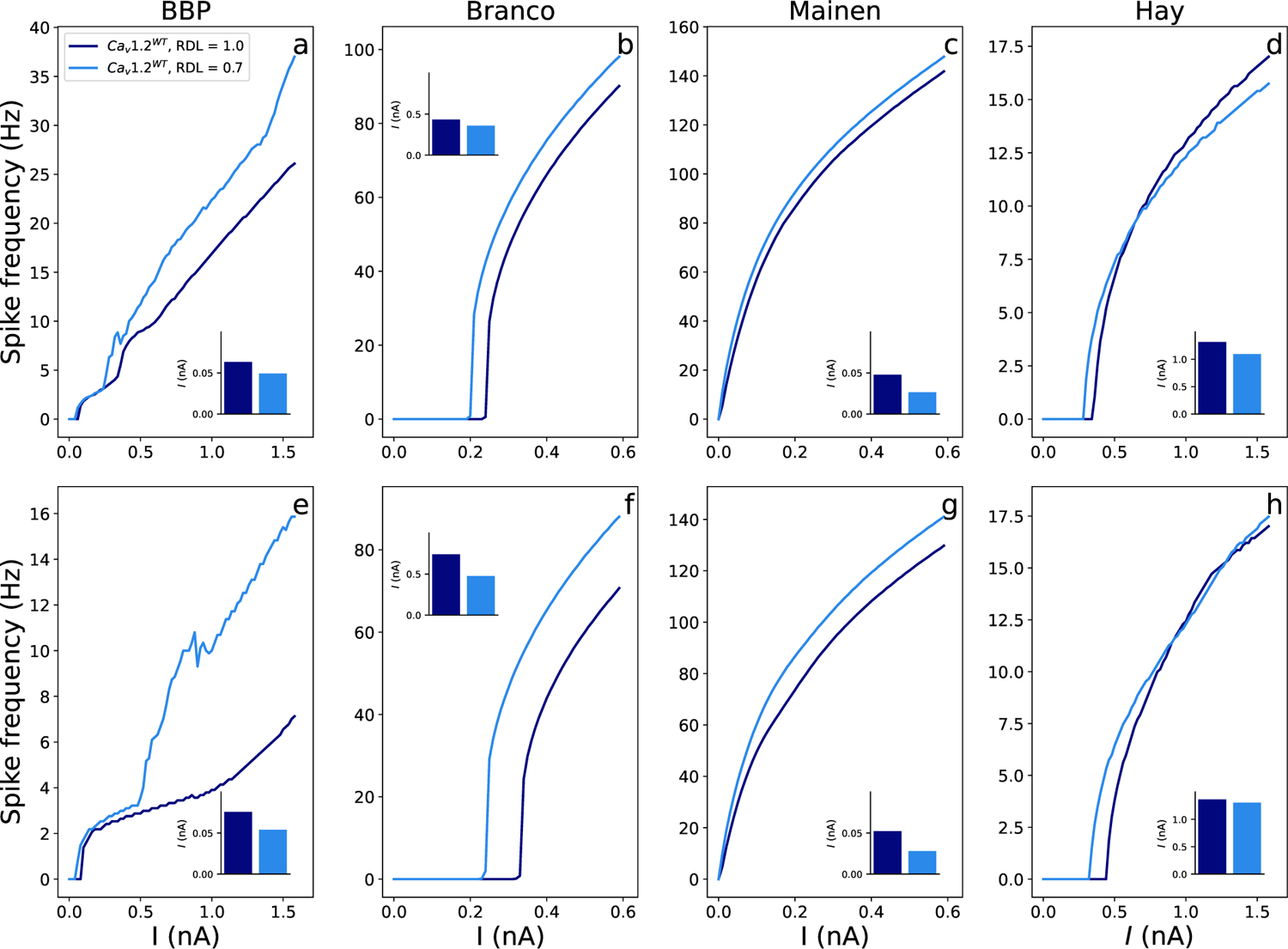
Effects of dendritic shortening on f-I curves and rheobase (inset axes) of multicompartmental L3PC models BBP (a,e), Branco (b,f), Mainen (c,g), and an L5PC Hay model (d,h). Upper panels (a-d) show somatic stimulation paradigm, and lower panels (e-h) show dendritic stimulation. RDL, relative dendritic length. For simulation protocol, see Figure 4.

Dendritic shortening also causes large drops in rheobase values in response to short-pulse stimulations (Figure 5 inset axes). For the somatic stimulation protocol, this is also due to reduction in the leak current to the reduced dendritic area. The heightened effect of shortening for apical stimulation protocols is likely a result of a far lower membrane area between stimulation and recording points, leading to a greater reduction in intracellular charge loss and heightened membrane responsiveness.

When the effects of morphological and electrophysiological variation are combined in BBP L23PCs during somatic stimulation, a slight rescuing effect occurs, particularly in the less extreme Ca_v_1.2^G402S^ variant (Figure 6a). This suggests that, until extremely high current inputs, shortened cells are more likely to display near-WT behaviours whereas as unshortened cells may show dampened excitability. However, when stimulated in the dendrite, the effect of shortening, though dampened, maintains a higher firing frequency at most current inputs (Figure 6e). In the alternative L23PC models, combined effects result in greater excitability, particularly in the Branco model, due to the extreme leftward shift resulting from dendrite shortening (Figure 6b-c,f-g). In the Branco and Hay models, the leftward shift in shortened cells is likely to alter cell input-output behaviour significantly (Figure 6b,d,f,h).

**Figure 6:**
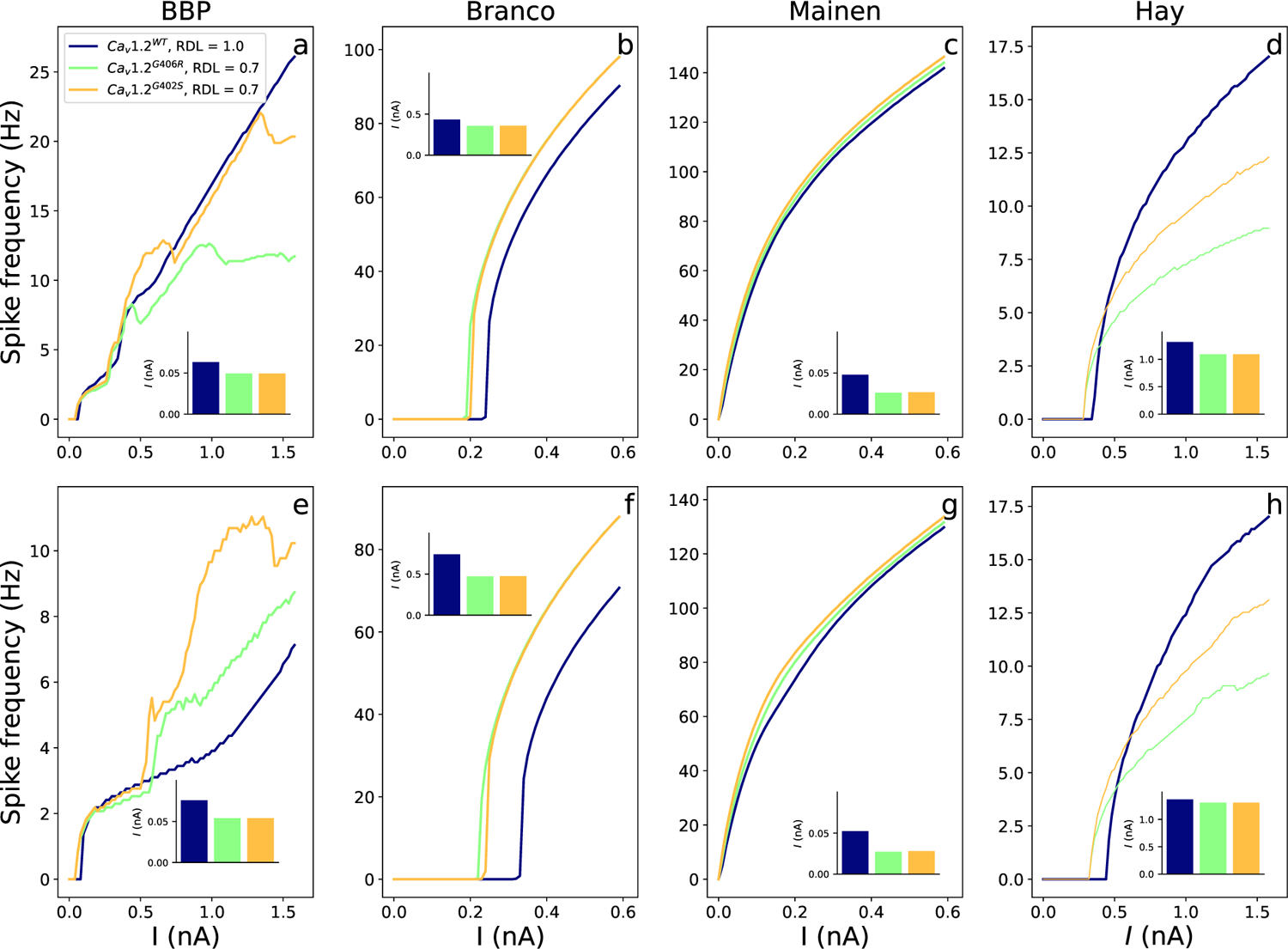
Combined effects of TS variant SSA/SSI alterations and dendritic shortening on f-I curves and rheobase (inset axes) of multicompartmental L3PC models BBP (a,e), Branco (b,f), Mainen (c,g), and an L5PC, Hay, model (d,h). Upper panels (a-d) show somatic stimulation paradigm, and lower panels (e-h) show dendritic stimulation. RDL, relative dendritic length. For simulation protocol, see Figure 4.

Shortened dendritic length is a key contributor to altered firing in TS, however, the effects of the SSA/SSI alterations somewhat compensate for this effect, with shortened TS neurones often showing a less extreme phenotype (Figure 4a-h, 6a-h).

Taken together, dendritic shortening in TS neurones markedly enhances cellular excitability by reducing dendritic leak, thereby lowering the firing threshold and increasing firing frequency, particularly during apical dendrite stimulation. This increased excitability is mitigated to some extent by the combined morphological and electrophysiological effects, leading to near-WT behaviours under somatic stimulation but maintaining higher firing frequencies under more biophysically relevant dendritic stimulation.

### TS Variants Alter Signal Integration in Pyramidal Cells

Spatiotemporal integration is critical to signal processing in neurones. We investigated the effects of TS variants on spatiotemporal integration in PCs by stimulating at the soma and the apical trunk at various intervals. This protocol employs two stimuli to simplify the typical pyramidal cell input: the apical trunk stimulus mimics synaptic input onto the apical dendrite, and the second, somatic, stimulus represents combined input from multiple basal dendritic branches. The BBP L23PC model shows no effect of TS variant SSA/SSI alterations on apical-somatic signal integration (Figure 7a). However, all three alternative models indicate that these variants will cause a widening of the signal integration curve (Figure 7b-d). In the Branco and Mainen L23PC models, only the Ca_v_1.2^G406R^ variant alters the curve from WT, suggesting the primary cause of this is the reduced voltage dependance of activation. The Hay model shows the greatest effect, which may be due cell-type differences or the strong I_CaHVA_ current in the “hot zone” of Ca^2+^ channels at the apical dendrite. The disparity between the BBP and alternative L23PC models likely arises due to the absence of dendritic I_CaHVA_ current in the BBP model.

**Figure 7:**
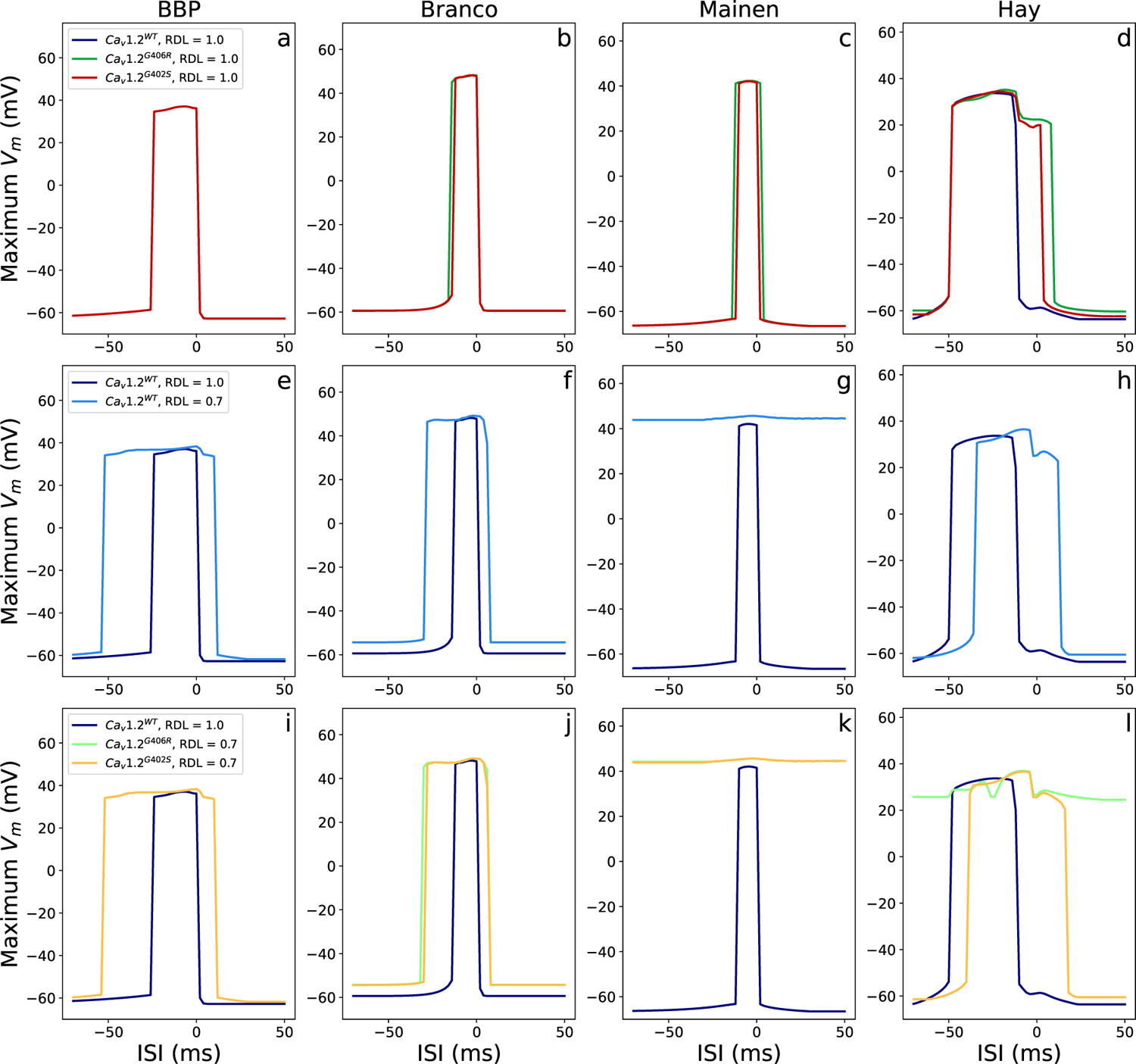
Independent effects of TS variant SSA/SSI alterations (a-d) cell shortening (e-h) and in combination (i-l) on the apical-somatic stimulus integration of multicompartmental L3PC models BBP (a,e,i), Branco (b,f,j), Mainen (c,g,k), and an L5PC, Hay, model (d,h,l). A modified interstimulus interval (ISI) protocol from Hay et al. [25] and Mäki-Marttunen et al. [53] was used to investigate stimulus integration in resting state neurones. Two stimuli were applied to the neurone; a square pulse (5ms) at the soma, and an EPSP pulse—emulating an excitatory postsynaptic potential—applied at a point 50% *−* 65% along the longest apical branch. The somatic stimulus was applied at 1000ms, and the order and interval between stimuli were dictated by the inter-stimulus interval, which ranged from *−*20–40ms. Somatic and apical input amplitudes for each model were: BBP 0.31nA; Branco 0.35nA; Mainen 0.025nA; Hay 0.8nA. RDL, relative dendritic length

All models show a large effect of dendrite shortening (Figure 7e-h). The Mainen model especially shows a high maximum membrane potential, even at large interstimulus intervals (Figure 7g). This suggests a single stimulus is sufficient to produce a strong response and the coincidence of a second stimulus is then inconsequential. This is also seen in the Hay model when variant SSA/SSI alterations are combined with dendritic shortening (Figure 7l).

Taken together, our results suggest the combined effects of SSA/SSI alterations and cell shortening are cumulative when I_CaHVA_ is present in the dendrite.

## 4 Discussion

This paper provides insights into the single-cell dynamics of TS-II cortical pyramidal neurones, investigating measures of excitability and signal integration. We utilise four distinct models to generate robust predictions and successfully delineate the potential impacts of morphological changes from the electrophysiological effects of this disorder. Although the outcomes of each model vary slightly, we find that alterations to neuronal activity generally concur with previous experimental results. Furthermore, the simulated effects of the morpho-logical changes described by Krey et al. [21] on neuronal responses were drastic and hitherto unstudied. In both cases, we provide testable predictions for *in vitro* analysis. Given the recent pre-clinical success of modulating TS-I exon expression in human induced pluripotent stem cell-derived neurones [22], these predictions inform experimental designs to further validate the electrophysiological impacts of TS-II variants and investigate potential therapeutic strategies. Similar previous *in silico* studies on the electrophysiological implications of TS have rightly focused on the myocardium, due to the high childhood mortality through cardiac events [16, 56]. However, the unique position of TS as a monogenic disorder causing autism spectrum disorder, and the strong links of Ca_v_1.2 with a plethora of neuropsychiatric disorders, creates a valuable opportunity to study the mechanisms of such disorders in the brain.

The altered gating kinetics of both TS-II channel variants enhanced the Ca^2+^ response to stimulus (Figure 3c,e). As expected, Ca_v_1.2^G406R^ showed a greater Ca^2+^ response than Ca_v_1.2^G402S^ [16, 17, 47]. This increased response lead to a reduction in firing frequency in three of four models, likely induced by the subsequent activation of SK channels having a hyperpolarising effect. The reduction in firing frequency is reflected in the flattened f-I curves of the BBP, Mainen, and Hay models (Figure 4a,c-e,g-h). Work with pharmacological L-type Ca^2+^ channel agonist, Bay-K, ratifies these findings in amygdala principal neurones [57], although antagonism with isradipine has also been shown to reduce spiking activity [58]. A Ca_v_1.2 forebrain knockout (Ca_v_1.2*^−/−^*) produces an opposite effect on the excitation of hippocampal PCs, despite a marginal, non-significant increase in the AHP [59]. Research in other cell types have found cell excitability is most strongly affected by Ca*_v_*1.3*^−/−^*, rather than Ca_v_1.2*^−/−^* [60], suggesting effects depend on the relative expression of the two channels in different cell types. Experimental results generally agree with our findings that Ca_v_1.2 activity does not impact the short pulse rheobase in TS-iPSCs [61] or other cell types [57, 59]. This invariability is likely due to the slow activation of Ca_v_1.2 channels and a delayed SK current response [62]. However, Lacinova et al. [58] found that pharmacological L-type Ca^2+^ channel agonism caused the threshold for long-stimulus evoked repetitive firing to decrease, as is seen in the frequency-current curve of the Ca_v_1.2^G406R^ variant in the Branco model.

This variant activates at a lower voltage and the reduced threshold response in this model, coupled with a lack of f-I flattening, may be indicative of a lower SK current response. SK channels are known to couple with L-type Ca^2+^ channels in the soma to regulate excitability [63].

Beyond its direct effects on cellular excitability, Ca^2+^ influx and the activation of Ca_v_1.2 pathways play a multitude of roles. This includes the modulation of dendritic extension by prolonging the duration of activity-dependent retraction events in the basal dendrites [21]. According to Krey et al. [21], the TS variant, Ca_v_1.2^G406R^, reduced the length of the dendrite by an average of 30% in human induced pluripotent stem cell-derived neurones and rat L23PCs. The proposed mechanism for this effect was a reduction in the binding efficiency between Ca_v_1.2^G406R^ and a GTP-binding protein, Gem [21]. Gem is known to inhibit both voltage-gated Ca^2+^ channels and key effectors involved in dendritic retraction [21, 64], although the relative contribution of these mechanisms to the observed effects remains unclear [21]. It is also unclear whether a similar binding disruption occurs to the same extent, or at all, with Ca_v_1.2^G402S^ channels, although the proximity of the *G*406 and *G*402 mutation sites suggests that this is possible. If such a disruption occurs, it may be that the higher voltage of activation of Ca_v_1.2^G402S^, and lower open probability, will result in a milder morphological phenotype. In the absence of this experimental data, we modelled a 30% reduction in basal and apical dendritic compartments for each variant. To verify our results, a comprehensive analysis of dendrite morphology and electrophysiological outcomes in neurones of both TS-II genotypes is required.

All models consistently predict that dendritic shortening increases cell excitability, as evidenced by changes in the f-I curve and rheobase (Figure 5). However, the predicted magnitude of the increase varies considerably by model. In most models, these effects are more pronounced when the stimulation is applied at the apical dendrite (Figure 5e,f,g). The dendritic stimulation protocol more closely follows *in vivo* conditions, suggesting TS PCs would be severely affected by shortening. Dendritic degeneration, modelled as shortening, has previously been functionally linked to neuronal hyperexcitability [65], and L3PCs have been shown to have reduced volume in human schizophrenia and autism spectrum disorder cohorts [66, 67]. Our predictions on the cellular impacts of dendritic shortening, in conjunction with these empirical findings, suggest that it could be a key mechanism underlying network dysregulation in neurodevelopmental and neuropsychiatric disorders.

When the models consider electrophysiological and morphological effects combined, the predicted outcomes differ. In the BBP and Hay models, the increase in excitation from dendritic shortening after somal stimulation is generally insufficient to raise the f-I curve above wild-type levels (Figure 6a,d), which is not the case for the Branco and Mainen models (Figure 6b,c). Due to the greater impact of dendritic shortening on excitability after apical stimulation, all cell models show more extreme phenotypes with dendritic stimulation (Figure 6e-h). Revah et al. [68] compared the cellular properties of stem cell-derived cortical organoids transplanted into the rat somatosensory cortex in both control and TS-I subjects. Transcriptomic mapping confirmed that these cells spanned all cortical layers, while their electrophysiological properties resembled human L23PCs. Notably, these neurones had reduced dendritic length and exhibited a lower maximum firing frequency when assessed via somatic whole cell patch-clamp. This aligns with predictions from the BBP and Hay models (Figure 6a,d), indicating that these more complex models best capture experimental results. Interestingly, the two models diverge in their predictions when stimulation occurs dendritically (Figure 6e,h). This discrepancy may arise from intrinsic morpho-electrical differences between the cell types or from the lack of dendritic I_CaHVA_ conductance in the BBP model. How dendritic shortening and TS variant electrophysiological changes interact *in vivo*, and whether such extreme cell type differences exist, remains in question. To our knowledge, no studies have experimentally described in detail how these changes interact in TS neurones, highlighting the need for further research to better understand the underlying mechanisms across the cortex.

Spatio-temporal signal processing is essential for the integration of multiple stimuli into a single output signal. In contrast to the BBP model, all alternative models predict that changes to Ca_v_1.2 SSA/SSI will influence apical-somatic signal integration in the neurone (Figure 7a-d). This discrepancy likely arises because the BBP model lacks I_CaHVA_ expression in the dendritic compartments, and therefore Ca_v_1.2 activation has no immediate effect on the response to an apical stimulus. While there is no L23PC specific Ca_v_1.2 distribution data available, studies in hippocampal PCs indicate high expression of Ca_v_1.2 in proximal dendrites and distal dendritic spines [29–31]. This suggests that the potential for dendritic I_CaHVA_ should not be overlooked. All alternative models predict a widening of the interstimulus interval area at which the two stimuli are integrated, which indicates a change in temporal coding and aligns with previous modelling work [53]. Furthermore, the Mainen and Hay models show the variants may cause a loss of order detection, as WT cells show absolute order detection, whereas shortened neurones can generate an output signal at positive interstimulus intervals. All models show a similar loss of order detection when dendritic shortening is implemented (Figure 7e-h). The Mainen model, however, predicts a more extreme alteration, with all stimulus protocols producing output in mutant neurones, regardless of the inter-stimulus interval (Figure 7k). This extreme effect is also predicted to occur in L5PCs in the Hay model, when dendritic shortening is combined with the SSA/SSI alterations of the Ca_v_1.2^G406R^ variant (Figure 7l). Single-cell electrophysiological data from TS PCs is needed to confirm our predictions that these neurones will show altered spatio-temporal signalling.

Dysregulation of excitability and input-output signalling in the single neurone is likely to have knock-on effects on cell-cell interactions through disruption to synaptic potentiation. In hippocampal pyramidal cells, a TS-II model conferring the Ca_v_1.2^G406R^ mutation showed enhanced excitatory post-synaptic potential amplitudes, reduced long-term potentiation induced by high-frequency stimuli and increased long-term potentiation induced by prolonged theta-train stimulation [69]. Although these are complex mechanisms, our findings may offer insights into part of the underlying processes. Higher excitatory post-synaptic potential responses, as seen in Sanderson et al. [69], could be underpinned by increased Ca^2+^ flux in the dendrites (Figure 3), resulting in increased potentiation responses under low-frequency theta-train stimuli. The changes in input-output signalling caused by shorter dendritic length (Figure 6) could further alter the induction of long-term potentiation *in vivo*. Potentiation and depression events also correlate to the structural remodelling of dendritic spines, which affects the number of synaptic connections possible. TS neurones develop a higher synaptic density than their WT counterparts [68]. Dendritic shortening is unlikely to affect intrinsic membrane excitability, as passive voltage responses are largely independent of dendritic length and branching patterns [70]. However, increased spine density in autism spectrum disorder and decreased spine density in schizophrenia [71–75], suggest that altered synaptic connectivity could be a mechanism of Ca^2+^-dependent disease aetiology. Hyperconnectivity in autism spectrum disorder could be a compensatory mechanism against dendritic shortening or result from altered single-cell excitability induced by heightened Ca^2+^ influx.

Understanding the complexities of how these disruptions at a single neurone level translate to broader network dysregulation is critical. Divergence of the frequency and pattern of spiking in shortened neurones may lead to a change in the signal strength ratio in local circuits relative to long-range connections. Increased Ca^2+^ current has been shown to have different effects on firing behaviour. In a TS-II Ca_v_1.2^G406R^ mouse model it was shown to promote burst over tonic firing [76] where Mäki-Marttunen et al. [77] found the opposite in L5PCs. Approximately 80% of L23PCs are thought to show tonic or intermediate firing phenotypes [78]. Unbalance in this ratio may have a significant influence on network communication, and notably changes in the burst and tonic firing balance have been related to several neuropsychiatric and neurodevelopmental disorders [79]. Further research is needed *in vitro* to determine the effects of TS mutations on spatio-temporal processing, how this may affect network activity, and whether this may form part of the pathology of the syndrome.

Our current study focuses on understanding post-developmental phenotypes, whereas many Ca_v_1.2-related effects may be established during development. Chen et al. [22] demonstrated that rescuing wild-type exon expression through targeted antisense oligonucleotide therapy can reverse morphological changes when applied during early post-natal development. However, this morphological rescue effect is unlikely to be as pronounced in later stages. Whether electrophysiological properties are similarly rescued, and the potential mitigating impact on phenotypic severity, remains unknown. By delineating the effects of morphology and electrophysiology on neuronal responses, our models suggest that partial rescue of electrophysiological properties, with limited rescue of morphological features, may improve f-I response profiles but will likely have minimal impact on signal integration in TS neurones. Further *in silico* network modeling, to elucidate the impact on network communication, could determine if these effects may justify investigating later-stage antisense oligonucleotide therapy.

Furthermore, increased Ca^2+^ influx through Ca_v_1.2 can also alter the induction of activity-dependent transcription [80, 81]. The full consequences of this are unknown but could be wide-ranging. Particularly in Ca_v_1.2^G406R^, the decreased VDA further decouples Ca_v_1.2 activation from spiking behaviour and activity-dependent transcription and this is thought to explain the increased neurological severity of TS-II Ca_v_1.2^G406R^ as compared to Ca_v_1.2^G402S^ [82]. We show that both TS variants—especially Ca_v_1.2^G406R^ —in conjunction with related morphological changes, can alter signal processing in PCs, which may disrupt activity-dependent transcription during critical developmental phases. Indeed, dysregulated gene expression in TS-II neurones during development results in aberrant cell differentiation, including a bias towards upper cortical layer cell types and a reduction in callosal projection neurones [61]— notably, reduced corpus callosum volume is strongly linked to autism spectrum disorder [83].

Autism spectrum disorder particularly is thought to arise from a developmental switch to favour local, over long-range, connectivity, and recent work has also linked this phenotype with a variety of neuropsychiatric disorders [84–86]. Similarly, the reduction in dendrite length modelled here may result in a disconnect between local and long-range signals. The impact of Ca_v_1.2^TSII^ variants on axonal retraction events remains unexplored, however, if the effect is similar to that in dendrites, it could significantly accentuate the bias towards local circuits. We, therefore, propose that while dysregulation of activity-dependent transcription is a significant factor in the neurological severity of Ca_v_1.2^G406R^ TS-II, other effects, particularly enhanced activity-dependent dendritic retraction and its coincidence with altered spiking behaviour, are also crucial in disease aetiology. Furthermore, these processes could influence the interconnectivity of brain regions, which has been posited as a theory for dysfunction in neurodevelopmental and neuropsychiatric disorders [85–87].

Several practical constraints impair the accuracy of these models. First and foremost is the availability of cell-type specific electrophysiological data for TS-II variants, as currently data is only available from heterologous models with widely different genetic backgrounds [16]. As an example, the shift in VDA caused by the Ca_v_1.2^G402S^ mutation has been reported at 2 or 10mV, depending on the cell model used [16, 88]. To ensure consistency, we simulated a +2mV shift, with all experimental data collected in the same heterologous model. Similarly, due to a lack of channel-specific expression and conductance data, we use a combined HVA Ca^2+^ current in place of Ca_v_1.2-specific values. HVA Ca^2+^ current combines all high voltage-activated calcium channels, comprising all Ca*_v_*1 and Ca*_v_*2 subtypes [43]. It is unclear to what extent each of these channels contributes to the combined current. The L-type channel Ca*_v_*1.3 likely comprises approximately 11% of L-type channels in the brain, with its expression partially overlapping with Ca_v_1.2 [1, 44]. In isolation, this would suggest that the use of a combined HVA Ca^2+^ current leads to a slight overestimation of TS-II effects in the model. However, TS-II mutations *in vivo* may induce a bias in exon splicing toward E8 inclusion. TS-I E8a mutations promote E8a over E8 inclusion [22]. Panagiotakos et al. [89] suggest this is due to their occurring within a motif at the exon border that is part of a splice acceptor site, facilitating E8a acceptance. Since E8 shares this conserved border motif [88], a similar mechanism could occur for E8. In this case, our model likely underestimates the effect of TS-II variants. A further limitation of the modelled current is the absence of calcium-dependent inactivation, which also varies by TS-II variant. However, no studies have investigated this effect in neurones and experiments done in myocardium have shown mixed results [88, 90, 91]. Finally, the variation within TS patients signifies a complex interconnected genetic architecture that is difficult to predict. The extremely high splice variation in *CACNA1C* and associations with various subunit isoforms will affect channel membrane trafficking, kinetics, and conductance [92].

Single-cell models, such as those used in this study, provide a base unit of computation from which a more complex picture can be built. Future studies should work towards developing network level information for Timothy Syndrome variants which would provide broader predictions for the roles of Ca_v_1.2 in Timothy Syndrome and other Ca^2+^-related disorders. Morphological and developmental data can be combined to create an accurate picture of the ratio of cell types and their predicted connections. Finally, incorporating dendritic spines, synaptic connections and biochemically accurate models of plasticity dynamics will enable investigations of these effects on network excitability [93–96].

## Acknowledgements

The authors would like to acknowledge the following funding: Academy of Finland (330776, 336376, and 318879), and University of Oslo Convergence Environment (4MENT).

